# Novel epitopes of human monoclonal antibodies targeting the influenza virus N1 neuraminidase

**DOI:** 10.1101/2021.02.26.433142

**Authors:** Ericka Kirkpatrick Roubidoux, Meagan McMahon, Juan Manuel Carreño, Christina Capuano, Kaijun Jiang, Viviana Simon, Harm van Bakel, Patrick Wilson, Florian Krammer

**Affiliations:** Department of Microbiology, Icahn School of Medicine at Mount Sinai, New York, NY; Graduate School of Biomedical Sciences, Icahn School of Medicine at Mount Sinai, New York, NY; Global Health Emerging Pathogens Institute, Icahn School of Medicine at Mount Sinai, NY, USA; Division of Infectious Diseases, Department of Medicine, Icahn School of Medicine at Mount Sinai, New York, NY, USA; Department of Genetics and Genomic Sciences, Icahn School of Medicine at Mount Sinai, New York, New York, USA; Icahn Institute for Data Science and Genomic Technology, Icahn School of Medicine at Mount Sinai, New York, New York, USA; Department of Medicine, Section of Rheumatology, the Knapp Center for Lupus and Immunology, University of Chicago, Chicago IL

## Abstract

Influenza virus neuraminidase (NA) targeting antibodies are an independent correlate of protection against infection. Antibodies against the NA act by blocking enzymatic activity, preventing virus release and transmission. As we advance the development of improved influenza virus vaccines that incorporate standard amounts of NA antigen, it is important to identify the antigenic targets of human monoclonal antibodies (mAbs). Additionally, it is important to understand how escape from mAbs changes viral fitness. Here, we describe escape mutants generated by serial passage of A/Netherlands/602/2009 (H1N1) in the presence of human anti-N1 mAbs. We observed escape mutations on the N1 protein around the enzymatic site (S364N, N369T and R430Q) and also detected escape mutations located on the sides and bottom of the NA (N88D, N270D and Q313K/R). We found that a majority of escape mutant viruses had increased fitness *in vitro* but not *in vivo*. This work increases our understanding of how human antibody responses target the N1 protein.

**Importance:** As improved influenza virus vaccines are being developed, the influenza virus neuraminidase (NA) is becoming an important new target for immune responses. By identifying novel epitopes of anti-NA antibodies, we can improve vaccine design. Additionally, characterizing changes in viruses containing mutations in these epitopes aids in identifying effects of NA antigenic drift.

## Introduction

Influenza viruses cause seasonal epidemics and, occasionally, global pandemics, that lead to significant morbidity and mortality worldwide (1, 2). They are a member of the family *Orthomyxoviridae* and contain a segmented, negative sense RNA genome. Two of the genetic segments encode for the glycoproteins present on the viral surface, the hemagglutinin (HA) and the neuraminidase (NA) (3, 4). The HA of influenza viruses, which is responsible for receptor binding and viral entry, has been largely credited as the immunodominant target of the immune response after vaccination and natural infection (3-5). The NA acts as a sialidase, removing terminal sialic acids and allowing for viral egress and spread. To function properly, the NA must be present on the viral surface as a homo-tetramer (6-8).

Seasonal influenza virus vaccines are the first line of defense against infection (9). Typically, these vaccines are standardized based on the HA content but have varying NA content with unknown structural integrity (10, 11). In addition, seasonal vaccines can have varying effectiveness from 20% to 60% in a given year (12, 13). Low vaccine effectiveness can be largely attributed to antigenic variability of the HA vaccine component compared to circulating strains (14-17). It may be possible to improve seasonal vaccine effectiveness by including a standard amount of a second viral antigen, the NA (18, 19). It has recently become appreciated as an additional important target of anti-influenza virus immunity (18-21). During natural infection, antibodies targeting both the HA and the NA are produced, however NA antibodies are rarely detected after vaccination (10). NA specific antibodies have been demonstrated to prevent severe infections, restrict transmission and protect from lethal challenge in the mouse model (8, 22-27). These antibodies function as neuraminidase inhibitors by blocking the NA enzymatic site and preventing viral spread (10, 22).

Residues critical for NA inhibiting antibodies were first characterized using murine antibodies (28-30). The monoclonal antibody (mAb) CD6 was found to span the dimer interface, while other mAbs were found to only bind to a single monomer. Additional work has been ongoing to identify targets of human mAbs (10, 31-33). A majority of these residues can be attributed to the discovery of broadly-reactive NA mAbs that target the enzymatic site (33). Interestingly, few residues have been identified as targets of both human and murine mAbs. This emphasizes the importance of using human anti-N1 mAbs to define true antigenic sites on the N1 protein. The targets of several previously published mAbs have yet to be defined, leaving a gap in our understanding of human mAb epitopes. Here, we use a panel of these uncharacterized mAbs to determine additional N1 residues targeted by human anti-N1 mAbs. The mAbs used in this study were isolated from individuals that were naturally infected and have varying levels of cross-reactivity and NAI activity (10).

## Results

### Generation of N1 mAb escape mutant viruses

For epitope analysis, we chose a panel of N1 specific mAbs from a recently published study (10). Our panel consisted of 8 mAbs: EM-2E01, 1000-1D05, 1000-3B04, 1000-3B06, 1000-3C05, 294-16-009-A-1C02, 294-16-009-A-1D05 and 300-16-005-G-2A04. We also included a negative IgG control antibody, KL-1C12, which targets the Ebola virus glycoprotein (34). Each mAb’s neuraminidase inhibition activity (NAI), measured using an enzyme-linked lectin assay (ELLA), and neutralization activity, measured through a plaque reduction assay (PRNA), was first determined against the wild type A/Netherlands/602/2009 H1N1 strain. All mAbs, aside from 1000-3C05 and 294-16-009-A-1D05, had NAI activity **(Table 1)**. The mAb 300-16-005-G-2A04 did not have neutralization activity, and mAbs 1000-3C05, 294-16-009-A-1C02, 294-16-009-A-1D05 had low neutralization activity **(Table 1)**.

**Table 1:**
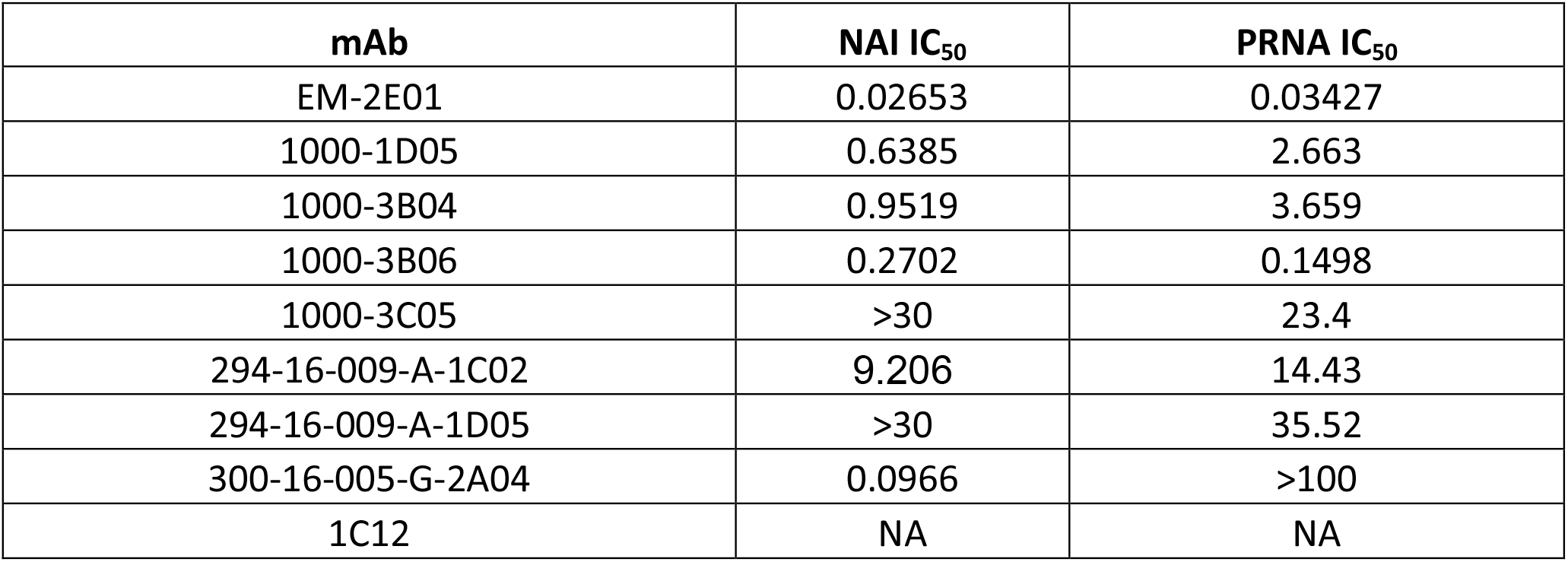
NAI and neutralization activities of mAbs against wild type virus. IC_50_ values are shown in µg/mL. NA stands for not applicable.

Escape mutant viruses (EMVs) were produced by passaging the wild type H1N1 virus with each antibody in Madin Darby Kidney cells (MDCKs). We began with a multiplicity of infection (MOI) of 0.01 and 0.25 times the 50% inhibitory concentration (IC_50_) of each mAb. EMVs were detected after 4-10 passages (2xIC_50_ to 128xIC_50_) **(Table 2)**. MAbs EM-2E01, 1000-3B04, 1000-3C05, 294-16-009-A-1C02 and 300-16-005-G-2A04 generated 7 distinct EMVs. The NA mutations identified were N88D (1000-3C05), N270D (1000-3B04), Q313K/R (294-16-009-A-1C02), S364N (EM-2E01), S364N/N369T (EM-2E01) and R430Q (300-16-005-G-2A04) **(Table 2)**. Additionally, the irrelevant IgG control virus contained an NA mutation at D454G. We were unable to generate viruses that escaped the mAbs 1000-1D05, 1000-3B06 and 294-16-009-A-1D05 after 10 passages. The detected mutations are distributed in different regions of the NA protein **(Figure 1)**. S364N, N369T and R430Q are located on the top of the tetramer **(Figure 1A)**. N270D and Q313K/R are located on the side of the tetramer **(Figure 1B-C)**. N88D is located on the bottom of the tetramer, near the head/stalk interface **(Figure 1C)** and R430Q is the closest to the NA enzymatic site. Mutations at N270 and N369 have also been identified using human mAbs in other studies (10, 31). The mutations N88D, Q313K/R, S364N and R430Q have not been previously identified using human mAbs. Each EMV and the irrelevant IgG control virus shared several HA mutations and also contained mutations in other genomic segments **(Tables 2)**.

**Table 2:**
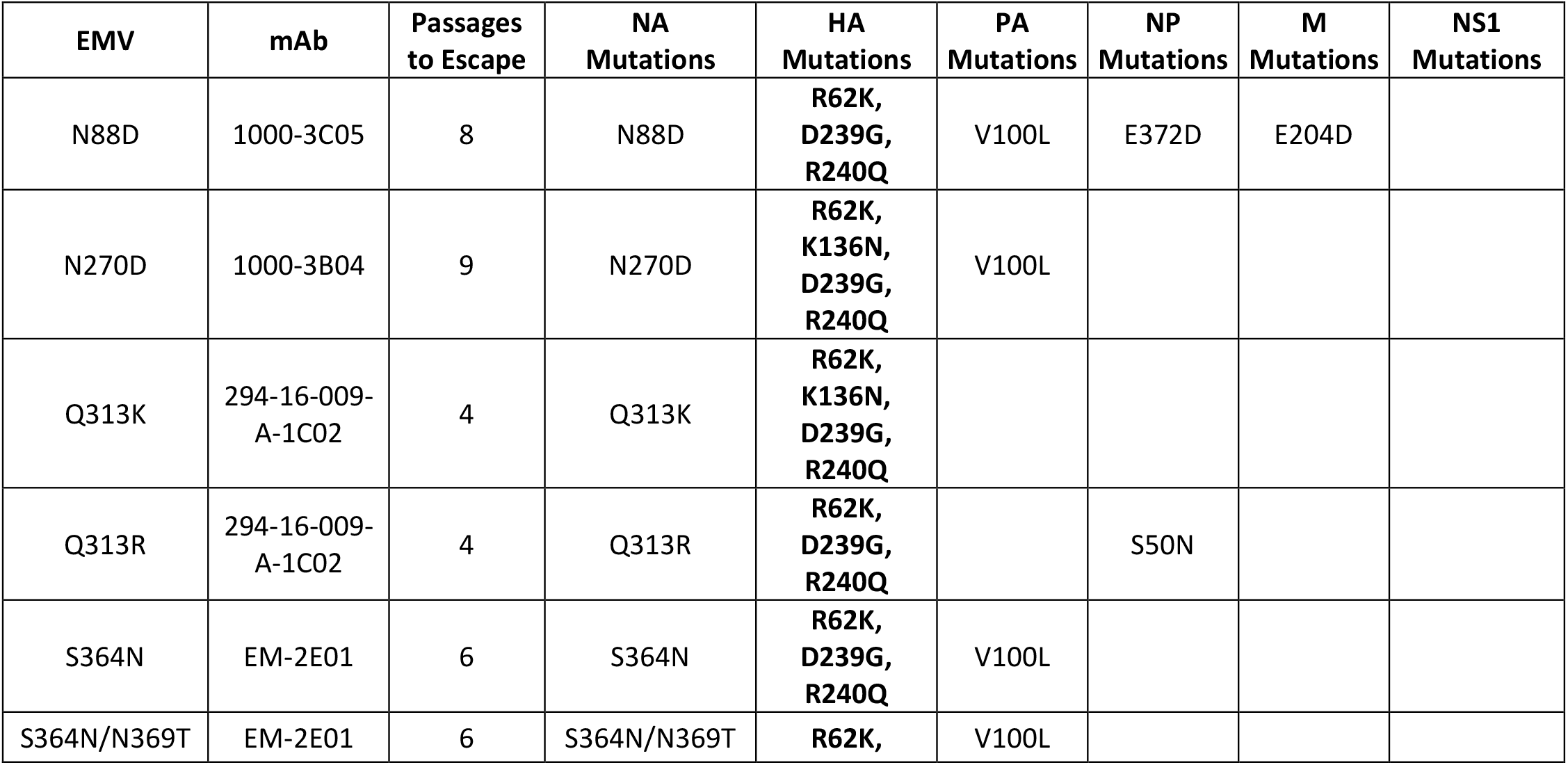
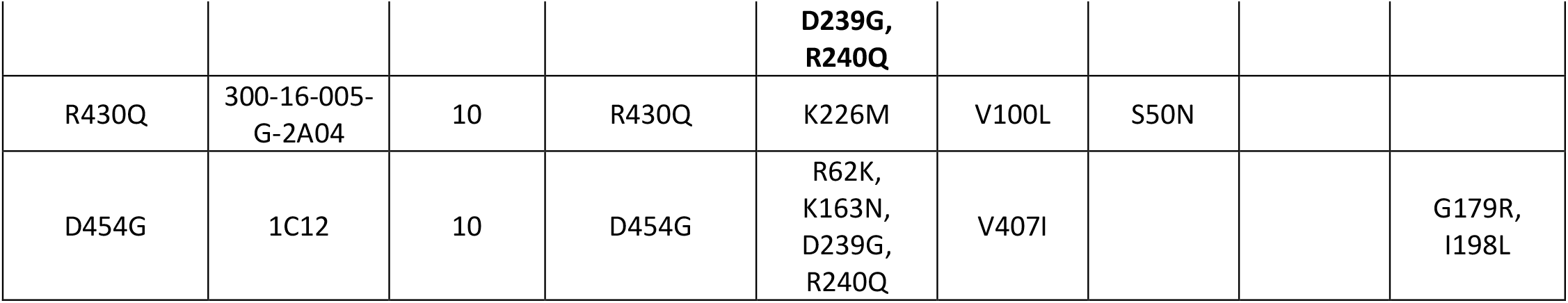
Escape mutations identified in passaged viruses. Specificity of each mAb was determined by referencing the previous publication characterizing these mAbs (10). Segments that are not listed did not contain any mutations. Numbering starts from the methionine of each protein. Mutations in bold are shared with the irrelevant IgG control virus.

**Figure 1:**
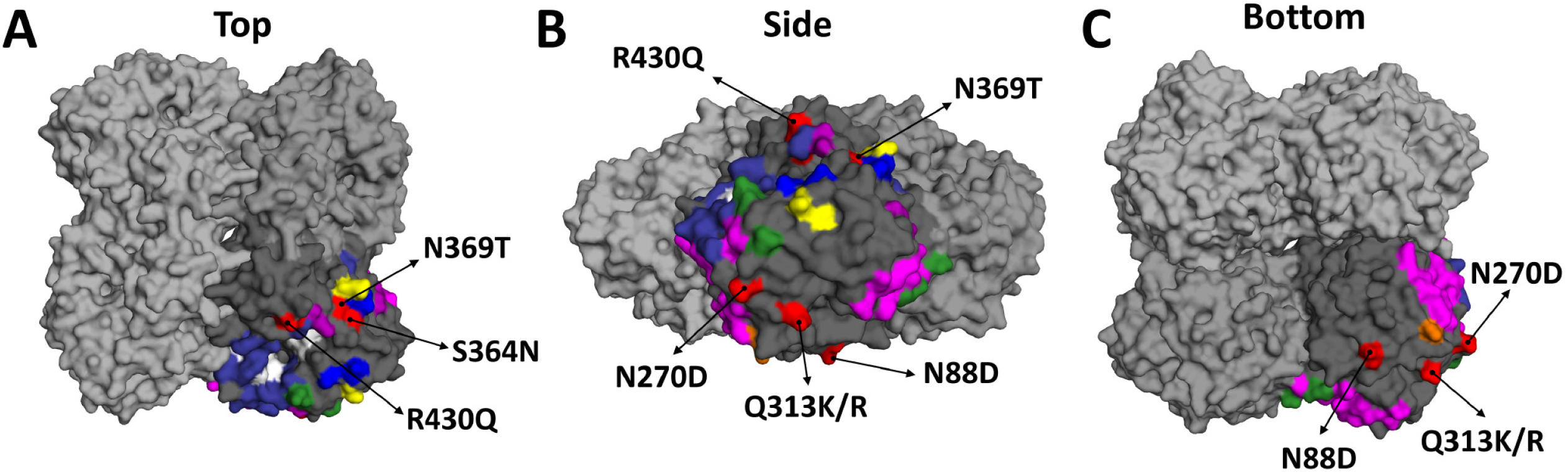
Escape mutations mapped onto a three-dimensional structure of the NA. The NA of A/California/04/2009 (PDB ID 3NSS(38)) is depicted as a tetramer with 3 monomers colored in light grey and one colored in darker grey. The darker grey subunit has residues identified by previous publications and the NA active site (in white). Murine epitopes are illustrated in blue (28), magenta (29) and yellow (30). Human epitopes are indicated in orange (10), purple (32), green (31) and indigo (33). Mutations identified in the EMVs used for this study are highlighted in red and identified using arrows. Views from the top **(A)**, side **(B)** and bottom **(C)** of the NA are depicted.

### Escape mutant viruses are resistant to binding, NAI and neutralization activity of mAbs

We next used the mAbs to evaluate the impact of each NA mutation on antibody binding, NAI and neutralization activity. Using immunofluorescence assays we identified that the N270D mutation impacted most of the antibodies in our panel, including those that produced other EMVs **(Figure 2A)**. The N88D mutation only affected the binding of the mAb that caused the mutation, 1000-3C05. Q313K impacted binding of the mAb that caused that mutation (294-16-009-A-1C02) and mAb 296-16-009-1D05 while Q313R only impacted the binding of 294-16-009-A-1C02. S364N and the double mutation S364N/N369T affected the binding of EM-2E01 and 1000-1D05 **(Figure 2A)**.

**Figure 2:**
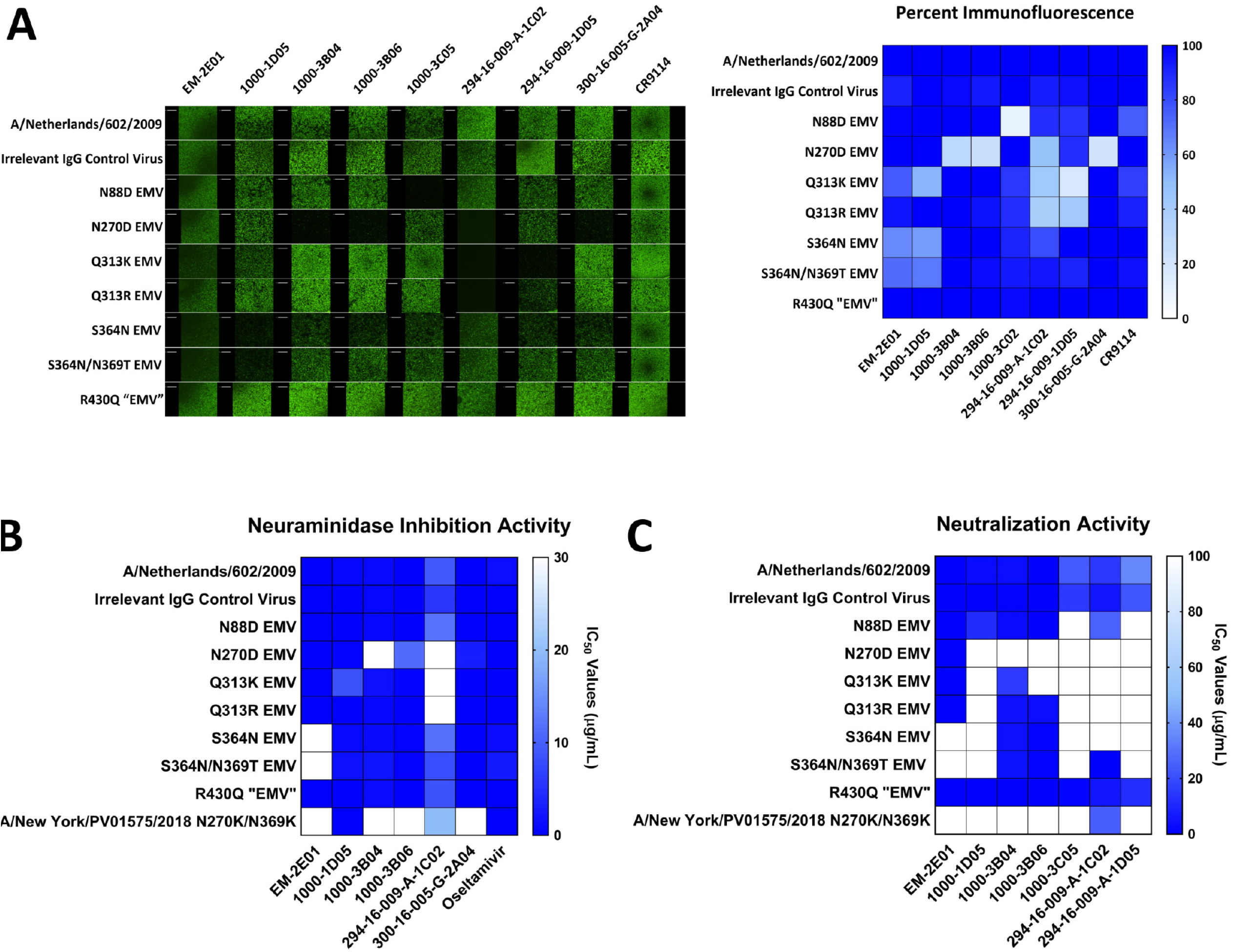
MAbs exhibit changes in binding, NAI and neutralizing activity towards EMVs. A) Immunofluorescence assay comparing binding of each mAb to wild type and EMVs. On the left are representative images and the right shows a heat map of percent luminescence compared to wild type. On the heat map, high binding is indicated by darker blue shading. Heat maps of NAI **(A)** and neutralization IC_50_s **(C)** of each mAb against wild type and EMVs. Each EMV is listed on the X-axis while mAbs are listed on the Y-axis. Darker blue is a higher IC_50_, which indicates stronger escape phenotypes. Binding, NAI and neutralization assays were conducted in duplicate.

To determine changes in NAI activity, we performed ELLAs with each EMV in the presence of each mAb. While 1000-3C05 has no NAI activity, we still assessed if the mutation it caused, N88D, impacted any other antibodies in the panel. We found that it caused no significant changes in the NAI activity of any of the mAbs tested **(Figure 2B)**. We found that the N270D mutation increased the NAI IC_50_ values of the same mAbs that lost binding to the EMV, including complete escape from 1000-3B04 and 294-16-009-A-1C02, along with resistance to 1000-3B06 (42-fold increase in NAI IC_50_) and 300-16-005-G-2A04 (30-fold change in NAI IC_50_) **(Figure 2B, Table 3)**. The Q313K mutation impacted the NAI activity of 1000-1D05 (13-fold increase in NAI IC_50_) and caused complete escape from 294-16-009-A-1C02. Q313R had less of an impact on mAb NAI activity and caused complete escape from 294-16-009-A-1C02 without causing resistance to other mAbs. S364N and the double S364N/N369T mutant led to complete escape from EM-2E01. The mutation R430Q did not have a strong impact on any mAb NAI activity **(Figure 2B)**. We also identified a natural isolate, A/New York/PV01575/2018, which contained mutations at residues identified in EMVs (N270K and N369K). This virus completely escaped EM-2E01, 1000-3B04, 1000-3B06 and 300-16-005-G-2A04 **(Figure 2B)**. No EMVs showed increased resistance to the neuraminidase inhibitor oseltamivir **(Figure 2B, Table 3)**. We observed a more significant impact on mAb neutralization compared to NAI activity. This may be caused by the mechanism of neutralization, which relies on strong NAI activity to prevent virus spread and plaque formation. So, moderate changes in NAI activity would still allow for the formation of plaques, increasing neutralizing IC_50_ values. The N88D EMV completely escaped mAbs 1000-3C05 and 300-16-005-G-2A04 **(Figure 2C, Table 4)**. The N270D EMV had complete escape from all mAbs in the panel aside from EM-2E01 **(Figure 2C, Table 4)**. Q313K led to escape from 1000-1D05, 1000-3B06, 1000-3C05, 294-16-009-A-1C02 and 300-16-005-G-2A04. However, Q313R led to escape from 1000-1D05, 1000-3C05, 294-16-009-A-1C02 and 300-16-005-G-2A04 but did not impact the neutralization activity of 1000-3B06. S364N and S364N/N369T EMVs completely escaped EM-2E01, 1000-1D05, 1000-3C05, 294-16-009-A-1C02 and 300-16-005-G-2A04 **(Figure 2C, Table 4)**. A/New York/PV01575/2018 escaped all mAbs in the panel aside from 294-16-009-A-1C02. We found that the R430Q “EMV” did not have resistance to any mAbs in the panel, preventing us from classifying it as a true escape mutant virus. Therefore, we do not think that R430Q is a critical residue for mAb escape, despite being tolerated in the NA. Importantly, the irrelevant IgG control virus had similar NAI and neutralization IC_50_ values compared to wild type virus, indicating that the mutation D454G does not directly impact mAb activity.

**Table 3:** NAI IC_50_ values for mAbs against all EMVs. IC_50_ values are shown in µg/mL. 30 µg/mL was the highest mAb concentration tested. Bolded values indicate significant differences in IC_50_s between the irrelevant IgG control virus and the EMV (*p<0.05).

| **EMV** | **EM-**<br>**2E01** | **1000-**<br>**1D05** | **1000-**<br>**3B04** | **1000-**<br>**3B06** | **294-16-009-**<br>**A-1C02** | **300-16-005 -**<br>**G-2A04** | **Oselta**<br>**mivir** |
| --- | --- | --- | --- | --- | --- | --- | --- |
| Irrelevant IgG Control Virus | 0.012 | 0.076 | 0.543 | 0.154 | 5.652 | 0.059 | 0.083 |
| N88D | 0.018 | 0.078 | 0.696 | 0.226 | 12.030 | 0.067 | 0.051 |
| N270D | 0.014 | 0.503 | **>30\*** | 11.250 | **>30\*** | 2.968 | 0.088 |
| Q313K | 0.032 | 8.502 | 1.663 | 0.453 | **>30\*** | 0.159 | 0.151 |
| Q313R | 0.015 | 0.113 | 0.774 | 0.190 | **>30\*** | 0.066 | 0.070 |
| S364N | **>30\*** | 0.898 | 1.017 | 0.320 | 11.770 | 0.125 | 1.249 |
| S364N/N369T | **>30\*** | 1.605 | 2.039 | 0.291 | 8.053 | 0.129 | 2.622 |
| R430Q | 0.019 | 0.140 | 1.054 | 0.242 | 8.563 | 0.107 | 0.056 |
| A/New<br>York/PV01575/2018<br>N270K/N369K | **>30\*** | 0.040 | **>30\*** | **>30\*** | 19.790 | **>30\*** | 0.064 |

**Table 4:** Neutralization IC_50_ values for mAbs against all EMVs. Values were determined using PRNAs. IC_50_ values are shown in µg/mL. 100 µg/mL was the highest mAb concentrated tested. Bolded values indicate significant differences in IC_50_s between the irrelevant IgG control virus and the EMV (*p<0.05).

| **EMV** | **EM-**<br>**2E01** | **1000-**<br>**1D05** | **1000-**<br>**3B04** | **1000-**<br>**3B06** | **1000-**<br>**3C05** | **294-16-009-**<br>**A-1C02** | **294-16-009-**<br>**A-1D05** |
| --- | --- | --- | --- | --- | --- | --- | --- |
| Irrelevant IgG Control Virus | 0.0300 | 0.341 | 0.8447 | 0.2041 | 14.98 | 5.592 | 22.34 |
| N88D | 0.0866 | 11.4 | 2.646 | 0.4076 | 100 | 25.56 | 100 |
| N270D | 0.0457 | **>100\*** | **>100\*** | **>100\*** | 100 | **>100\*** | 100 |
| Q313K | 0.2807 | **>100\*** | 15.21 | **>100\*** | 100 | **>100\*** | 100 |
| Q313R | 0.1557 | >100* | 4.322 | 3.477 | 100 | >100* | 100 |
| S364N | >100* | >100* | 4.098 | 1.196 | 100 | >100* | 100 |
| S364N/N369T | >100* | >100* | 4.504 | 0.8054 | 100 | >100* | 100 |
| R430Q | 0.0320 | 0.0706<br>9 | 0.3104 | 0.1467 | 1.216 | 5.683 | 12.03 |
| A/New<br>York/PV01575/2018<br>N270K/N369K | >100* | >100* | >100* | >100* | 100 | 24.4 | 100 |

### Escape mutant viruses retain fitness *in vitro* and *in vivo*

Evaluating the fitness of each EMV is important to determine if natural isolates that acquire these mutations will have a probability of showing increased or decreased fitness. We assessed growth kinetics in two cell lines, MDCKs and A549 cells (derived from human lung epithelial cells). In MDCK cells, EMVs aside from N270D had increased titers compared to the irrelevant IgG control virus, with peak titers being 10-fold higher at 48 hours post infection **(Figure 3A)**. This pattern was similar in A549 cells, with N270D and R430Q EMVs replicating similarly to the irrelevant IgG control viruses **(Figure 3C)**. We also calculated the area under the curve (AUC) for all EMVs. Using AUC values, we were able to determine that the N88D, Q313K, Q313R, S364N and S364N/N369T EMVs had significantly higher growth kinetics in MDCK cells **(Figure 3B)**. The AUC calculations in A549 cells indicated that the N88D, Q313K, Q313R and S364N/N369T EMVs also had significantly better growth kinetics in A549 cells **(Figure 3D)**.

**Figure 3:**
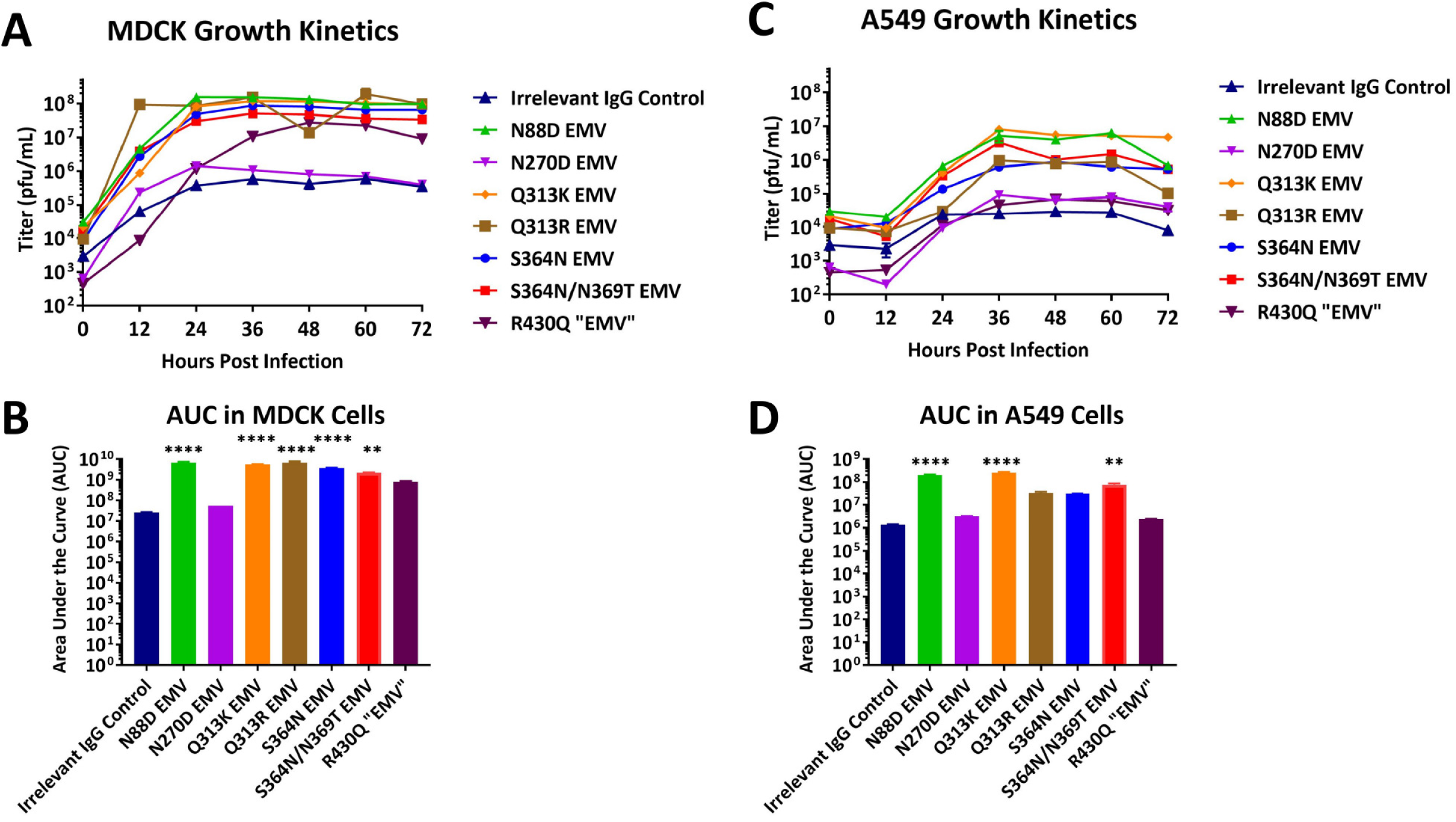
EMVs have an increased growth phenotype *in vitro*. EMVs’ growth curves and their corresponding AUC values are shown for MDCK **(A-B)** and A459 **(C-D)** cell lines. Samples were collected every 12 hours. Time 0 is a measurement of starting inoculum titers. The significance between AUC values for EMVs compared to the irrelevant IgG control virus are indicated on the figure (**p<0.01, ***p<0.001 and ****p<0.0001). Figures show the mean with standard deviations indicated by error bars, when applicable.

Additionally, we determined the mouse 50% lethal dose (mLD_50_) of each EMV to assess fitness changes *in vivo*. The irrelevant IgG control virus had a similar mLD_50_ to the wild type virus, 7 and 1 plaque forming units (pfu), respectively **(Table 5)**. This indicates that the irrelevant IgG control virus G454D mutation is not influencing fitness *in vivo*. Since both Q313R EMV and S364N EMV had similar changes in binding, NAI and neutralization activity to Q313K and S364N/N369T EMVs, only one was chosen for mLD_50_ experiments. A majority of EMVs had similar mLD_50_ values as the irrelevant IgG control virus including the N88D, N270D, Q313K and S364N/N369T EMVs **(Table 5)**. The R430Q EMV had a moderate (45-fold) increase in mLD_50_ **(Table 5)**. However, we also noted that this EMV contained a unique HA mutation, K226M, while the other EMVs shared 3-4 HA mutations with the irrelevant IgG control virus. This data suggests that mutations in the NA have less of an impact on viral fitness when compared to mutations in the HA stalk(35). Overall, it appears that mutations that have impacts on binding, NAI and neutralization activities may lead to increases of *in vitro* fitness however they do not significantly change fitness of the virus *in vivo* in the mouse model.

**Table 5:**
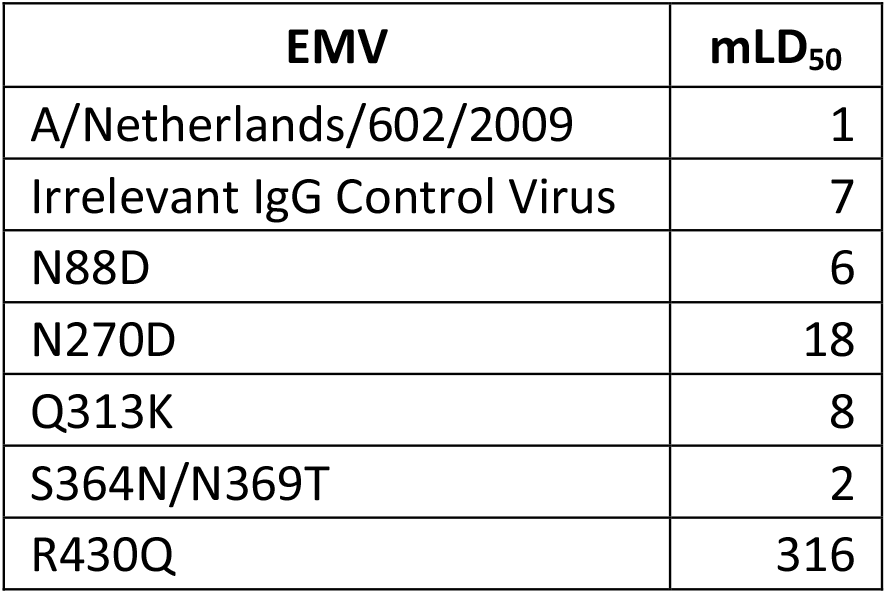
EMVs have similar mLD_50_ values as the irrelevant IgG control virus. Values are listed as pfu/mouse.

## Discussion

Our study has identified novel epitopes on the N1 targeted by human mAbs. Only 2 of the escape mutations detailed here have been previously reported, N270 and N369, indicating that these 2 residues are frequently targeted epitopes of human mAbs (10, 28, 31). Interestingly, a majority of 2017-2018 isolates contained N270K and N369K mutations, which further emphasizes their importance for mAb binding and NAI activity. The remaining mutations, N88D, Q313K/R, S364N and R430Q, are part of newly identified mAb epitopes.

The mutation N88D, a critical residue for 1000-3C05, is located very close to the NA head-stalk interface. MAb 1000-3C05 does not exhibit NAI activity and is poorly neutralizing, however, previous reports have noted that it is protective *in vivo*, cross-reactive with several human N1s (pre- and post-pandemic) and can utilize Fc-effector functions (10, 32). N88 is most likely not a direct target of 1000-3C05 based on its location on the NA and may induce an allosteric change to ablate mAb binding instead.

Mutations at Q313 were necessary for the evasion of 296-16-009-A-1C02 during escape mutagenesis. Once the EMV was identified, we noted that this residue was also important for another mAb in our panel, namely 1000-1D05. This residue is located on the side of the NA, outside of any previously defined antigenic regions. We noticed that the Q313K mutation had a slightly stronger effect on mAb escape compared to Q313R. This may be best explained by amino acid biochemistry. The mutation from glutamine to asparagine does not change side chain acidity/basicity however, lysine is a change from a neutral to a basic side chain.

We identified two separate EMVs containing the mutation S364N. When comparing escape phenotypes, the S364N and S364N/N369T EMVs completely escaped the mAb EM-2E01, became resistant to neutralization of 1000-1D05 and remained sensitive to the remaining mAbs in the panel. These data suggest that S364N is sufficient for escape from EM-2E01 and that N369T is not critical to mAb escape. Furthermore, the S364N mutation alone is responsible for the introduction of an N-linked glycosylation, which is likely responsible for blocking mAb activity. This is interesting because many recent isolates contain mutations at N369, like the A/New York/PV01575/2018 isolate used in this study, but S364 is highly conserved.

The mutation G454D was identified in the irrelevant IgG control virus. However, this virus behaved very similarly to wild type, indicating that this mutation does not significantly impact viral fitness. To truly understand how these mutations impact NA antigenicity, it would be important for future studies to test how human sera inhibit the neuraminidase activity of our EMVs.

When combined with previous reports, we can conclude that human antibodies are targeting more than just the enzymatic site (10, 31-33). **Figure 1** illustrates where the EMV mutations, along with others discussed here, are located on the NA. Aside from overlaps at 248, 249, 270, 273, 309, 369, 451 and 456, the epitopes for murine mAbs are unique compared to what has been observed for human mAbs. This highlights why it is important to evaluate antigenic sites using human monoclonal antibodies to increase understanding of how the N1 is being targeted by our immune responses.

## Methods

### Cells, virus and antibodies

MDCK cells (ATCC #CCL-34) and A549 cells (ATCC #CCL-185) were obtained from American Type Culture Collection (ATCC) and propagated using cDMEM (1x Dulbecco’s Modified Eagle Medium [Gibco], 10% heat inactivated fetal bovine serum [Sigma-Aldrich], 1U/mL penicillin-1µg/mL streptomycin solution [Gibco] and 10mM 4-(2-hydroxyethyl)-1-piperazineethanesulfonic acid [HEPES, Gibco]).

A/Netherlands/602/2009 (pdmH1N1) was grown in our laboratory by injection of 10-day old specific pathogen free (SPF) embryonated chicken eggs (Charles River Laboratories) at 37°C for 2 days. All mAbs were identified and isolated previously and provided by Dr. Patrick Wilson (University of Chicago) (10). They were expressed in our lab using the Expi293 transfection kit according to the manufacturer’s instructions (ThermoFisher). MAbs were purified through gravity flow with protein G sepharose packed columns and concentrated as described previously (36).

### Escape mutant generation

Escape mutant viruses were generated using the H1N1 virus A/Netherlands/602/2009 as maternal strain. MDCK cells (ATCC CCL-34) were plated at 6×10^5^ cells/mL in a 12-well, sterile cell culture plate and incubated overnight at 37°C with 5% CO_2_. The following day, virus was diluted to an MOI of 0.01 (1×10^3^ pfu) in 1x minimal essential medium (1xMEM; 10% 10xMEM [Gibco], 2mM L-glutamine [Gibco], 0.1% sodium bicarbonate [Gibco], 10 mM HEPES, 1U/mL penicillin-1µg/mL streptomycin solution, and 0.2% bovine serum albumin) supplemented with 1µg/mL tolylsulfonyl phenylalanyl chloromethyl ketone (TPCK)-treated trypsin. Antibodies were then added to the virus at a concentration that was 0.25 times the NAI IC_50_. The virus and antibody mixture was incubated for 1 hour at room temperature, shaking. MDCK cells were washed with 1x phosphate buffered saline (1xPBS, Gibco) and then the mixture was added to the cells. They were then incubated at 37°C with 5% CO_2_ for 2 days. Supernatant was collected stored at -80°C until further use. For subsequent passages, MDCK cells were plated as described above and infected with a 1:10 dilution of the previous passage in 1xMEM with TPCK-treated trypsin (1µg/mL) and incubated for 40 minutes at 37°C with 5% CO_2_. In the meantime, mAb was diluted in 1xMEM with TPCK-treated trypsin to a concentration that was doubled from the previous passage (0.25x, 0.5x, 1x and so on). After 40 minutes, diluted mAb was added to the virus infected cells and left for 2 days at 37°C with 5% CO_2_. Cell culture supernatant was screened for escape mutant viruses through plaque assays with 128xIC_50_ of mAb present in the agarose overlay. Individual plaques were chosen and propagated in SPF eggs for 2 days at 37°C as described above.

### RNA isolation and deep sequencing

RNA was isolated from egg allantoic fluid using the E.Z.N.A viral RNA extraction kit (Omega Bio-Tek) according to the manufacturer’s instructions and then underwent next generation sequencing. Sequences were assembled using a pipeline designed at the Icahn School of Medicine at Mount Sinai as described previously (37). To identify point mutations, full length coding sequences were compared to the sequenced wild type A/Netherlands/602/2009 used for escape mutagenesis.

### Plaque assay

Plaque assays were performed using a standard protocol. MDCK cells were seeded 24 hours prior at 8×10^5^ cells/mL. Next, virus samples were serially diluted in 1xMEM from 10^−1^ to 10^−6^. MDCK cells were washed with 1xPBS and then infected with 200µL of each virus dilution. Virus was incubated for 40 minutes at 37°C with 5% CO_2_, rocking every 10 minutes. Afterwards, virus was aspirated and immediately replaced with 1mL of an agarose overlay containing 2xMEM, 0.1% (diethylaminoethyl)-dextran (DEAE), 1µg/mL TPCK-treated trypsin, and 0.64% Oxoid agarose. Plates were incubated for 2 days at 37°C with 5% CO_2_. Cells were then fixed using a 3.7% solution of paraformaldehyde (PFA) and incubated at 4°C overnight. To plaque visualization, the overlay was removed and cells were stained with a solution containing 20% methanol and 0.5% crystal violet.

### Immunofluorescence

MDCK cells were plated at 3×10^5^ cells/mL in a sterile, 96-well plate and incubated overnight at 37°C with 5% CO_2_. The following day, cells were checked for >99% confluency and washed with 1xPBS. Virus was diluted to an MOI of 5 in 1xMEM and added to each well (100µL/well). Plates were incubated overnight at 37°C with 5% CO_2_. The following day, cells were fixed using 200µL of 3.7% PFA and incubated overnight at 4°C. Next, the PFA was removed and cells were blocked with 1xPBS containing 3% nonfat milk (American Bio) for 1 hour at room temperature. The blocking solution was then removed and replaced 1% nonfat milk. Primary mAbs were diluted to 300µg in 1xPBS and added to the 1% milk at a 1:10 dilution, for a final concentration of 30µg per well. Primary antibodies were incubated for 1 hour at room temperature, shaking. Plates were then washed 3 times with 1xPBS. Secondary antibody AlexaFluor™ 488 goat anti-human IgG (H+L) (Invitrogen) was diluted to 1:500 in 1% milk, added and incubated for 1 hour at room temperature, in the dark and shaking. The plates were then washed 3 times with 1xPBS. To prevent cells from drying out, 50µL of 1xPBS was added to each well. Plates were visualized using the CELIGO S adherent cell cytometer (Nexcelom Bioscience) with the 2-channel Target 1+2 (merge) setting. Exposure time, gain and focus (set using image-based auto focus with the 488nm signal as the target) were automatically determined by the machine. Fluorescence was calculated using the default analysis settings and percent florescence was determined based on wild type signal. We performed 2 independent assays, however only representative images from one assay are shown here.

### Enzyme-linked lectin assay (ELLA)

We performed enzyme-linked lectin assays with each EMV and wild type virus to determine NA activity. Flat bottom Immulon 4HBX microtiter plates (Thermo Scientific) were coated with 25µg/mL of fetuin (Sigma) diluted in 1xPBS, at 100µL per well, and incubated overnight at 4°C. The next day, viruses were serially diluted (3-fold) in sample diluent buffer (1xPBS with 0.5mM MgCl_2_, 0.9mM CaCl_2_, 1% BSA and 0.5% Tween 20) in a sterile 96-well plate. Once diluted, an additional 1:1 ratio of sample diluent was added to the plate. This was incubated for 1 hour at room temperature, shaking. After 1 hour, the fetuin coated plates were washed 3 times with PBS containing 0.1% Tween 20 (PBS-T) using the AquaMax 3000 automated plate washer. Diluted virus was immediately added to the plates and then incubated at 37°C, with 5% CO_2_ for 18 hours (overnight). The following day, plates were washed 6 times with PBS-T. Peanut agglutinin (PNA, Sigma) was diluted to 5µg/mL in conjugate diluent buffer (1xPBS with 0.5mM MgCl_2_, 0.9mM CaCl_2_ and 1% BSA) and added to the washed plates. This was incubated for 2 hours in the dark at room temperature. The PNA was then removed and plates were washed 3 times with PBS-T. SigmaFast *o*-phenylenediamine dihydrochloride (OPD, Sigma) was diluted in water. OPD was added at 100µL per well and incubated for 3 minutes at room temperature. The reaction was stopped by adding 50µL of 3M hydrochloric acid and then absorbance (at 490nm) was immediately determined using Synergy H1 hybrid multimode microplate reader (BioTek). PRISM 7.0 was used to determine the effective concentration of each virus that would yield detectable NA activity. Each ELLA was done in triplicate.

### Neuraminidase inhibition assay (NAI assay)

To determine NAI activity of each mAb, flat-bottom Immulon 4HBX microtiter plates (Thermo Scientific) were coated with25µg/mL fetuin (Sigma) diluted in 1xPBS and incubated overnight at 4°C. The following day, antibodies were diluted in sample diluent buffer to 120µg/mL and then serially diluted 1:3 in a sterile 96-well plate. Virus was diluted to 2 times the effective concentration determined in an ELLA and added to mAbs at a 1:1 ratio. This was incubated for 1 hour at room temperature, shaking. The fetuin coated plates were washed 3 times in PBS-T as described above. Virus-mAb dilutions were transferred to the fetuin coated plates and incubated at 37°C, with 5% CO_2_, for 18 hours (overnight). The following day, we followed the ELLA procedure described above. The 50% inhibitory concentration (IC_50_) was determined using PRISM 7.0. Each NAI assay was performed in duplicate. Significance between IC_50_ values for each EMV and the irrelevant IgG control virus were calculated using a 2-way ANOVA.

### Plaque reduction assay (PRNA)

MDCK cells were plated at 8×10^5^ cells/mL in 12-well plates. The following day, mAbs were diluted to 100µg/mL in 300µL 1xMEM and serially diluted 1:5 in 1xMEM to a final concentration of 0.032µg/mL. Each virus was then diluted in 1xMEM to 1×10^3^ pfu and 50µL was added to each antibody dilution. This virus-mAb mixture was incubated for 1 hour at room temperature, shaking. Afterwards, MDCK cells were washed with 1xPBS and then immediately infected with 200µL per well of the virus-mAb mixture. The plates were incubated for 40 minutes at 37°C, with 5% CO_2_, rocking every 10 minutes. In the meantime, the overlay was prepared. Antibodies were diluted to 100µg/mL in 625µL of 2xMEM and then serially diluted as described above. TA solution containing 1xDEAE and 1µg/mL of TPCK-treated trypsin in sterile water for injection (Gibco) was added at 180µL to each antibody dilution. When the infection finished, 360µL of 2% Oxoid agarose was added the overlay mixture, in small batches to prevent solidification before being transferred to cells. The inoculum was removed and immediately replaced with the overlay so that the mAb dilution in the overlay was the same as the concentration in the inoculum. The plates were then incubated at 37°C, with 5% CO_2_, for 2 days. Cells were fixed and stained as described above. Each PRNA was done in duplicate. Significance between neutralizing IC_50_ values for each EMV and the irrelevant IgG control virus were calculated using a 2-way ANOVA.

### Growth kinetics

MDCK cells were seeded at 8×10^5^ cells/mL in sterile, 24-well plates and incubated overnight at 37°C with 5% CO_2_. The following day, virus was diluted to an MOI of 0.01 (5×10^3^ pfu) in 1xMEM supplemented with 0.2µg/mL of TPCK-treated trypsin. Cells were washed with 1xPBS and then immediately infected. A portion of the inoculum was reserved as the T0 control. The plates were incubated for 72 hours and an aliquot was collected every 12 hours, for 72 hours total. Virus titers were determined using plaque assays. Each growth curve was performed in biological duplicates. AUC values were calculated using PRISM 7.0. Significance was determined using a one-way ANOVA with the default PRISM settings.

### Mouse lethal dose (mLD_50_)

The mLD_50_ for each EMV was determined using female BALB/c mice (at 6-8 weeks of age, Jackson Laboratory) in accordance with protocols approved by the Institutional Animal Care and Use Committee at the Icahn School of Medicine at Mount Sinai. Each virus was diluted from 10^5^ to 10^1^ in 1xPBS and 3 mice per dilution were infected (50µL per mouse). Weight loss and survival was monitored daily, for 14 days post infection. Mice that lost more than 25% of their initial body weight were euthanized.

## Acknowledgements

We thank the Mount Sinai Pathogen Surveillance Program for providing access to discard nasopharyngeal swab specimen. This work was supported in part by the National Institute of Allergy and Infectious Disease (NIAID) Centers of Excellence for Influenza Research and Surveillance (CEIRS) contract (HHSN272201400008C and HHSN272201400005C).

## References

1. Prevention CfDCa. 2020. Past Seasons Estimated Influenza Disease Burden. https://www.cdc.gov/flu/about/burden/past-seasons.html.Accessed

2. Thompson WW, Weintraub E, Dhankhar P, Cheng PY, Brammer L, Meltzer MI, Bresee JS, Shay DK. 2009. Estimates of US influenza-associated deaths made using four different methods. Influenza Other Respir Viruses 3:37–49.

3. Shaw M, Palese P. 2013. Field’s Virology, p 1151–1185. In HP Knipe DM (ed), 6 ed. Lippincott-Raven, Philadelphia, PA.

4. Bouvier NM, Palese P. 2008. The biology of influenza viruses. Vaccine 26 Suppl 4:D49–53.

5. Nachbagauer R, Liu WC, Choi A, Wohlbold TJ, Atlas T, Rajendran M, Solórzano A, Berlanda-Scorza F, García-Sastre A, Palese P, Albrecht RA, Krammer F. 2017. A universal influenza virus vaccine candidate confers protection against pandemic H1N1 infection in preclinical ferret studies. NPJ Vaccines 2:26.

6. McAuley JL, Gilbertson BP, Trifkovic S, Brown LE, McKimm-Breschkin JL. 2019. Influenza Virus Neuraminidase Structure and Functions. Front Microbiol 10:39.

7. Air GM. 2012. Influenza neuraminidase. Influenza and Other Respiratory Viruses 6:245–256.

8. McMahon M, Kirkpatrick E, Stadlbauer D, Strohmeier S, Bouvier NM, Krammer F. 2019. Mucosal Immunity against Neuraminidase Prevents Influenza B Virus Transmission in Guinea Pigs. mBio 10:e00560–19.

9. Grohskopf LA, Alyanak E, Broder KR, Walter EB, Fry AM, Jernigan DB. 2019. Prevention and Control of Seasonal Influenza with Vaccines: Recommendations of the Advisory Committee on Immunization Practices — United States, 2019–20 Influenza Season. MMWR Recomm Rep 68:1–20.

10. Chen YQ, Wohlbold TJ, Zheng NY, Huang M, Huang Y, Neu KE, Lee J, Wan H, Rojas KT, Kirkpatrick E, Henry C, Palm AE, Stamper CT, Lan LY, Topham DJ, Treanor J, Wrammert J, Ahmed R, Eichelberger MC, Georgiou G, Krammer F, Wilson PC. 2018. Influenza Infection in Humans Induces Broadly Cross-Reactive and Protective Neuraminidase-Reactive Antibodies. Cell 173:417-429.e10.

11. Wohlbold TJ, Nachbagauer R, Xu H, Tan GS, Hirsh A, Brokstad KA, Cox RJ, Palese P, Krammer F. 2015. Vaccination with Adjuvanted Recombinant Neuraminidase Induces Broad Heterologous, but Not Heterosubtypic, Cross-Protection against Influenza Virus Infection in Mice. mBio 6:e02556–14.

12. Centers for Disease Control and Prevention NCfIaRDN. 2020. Past Seasons Vaccine Effectiveness Estimates. https://www.cdc.gov/flu/vaccines-work/past-seasons-estimates.html. xAccessed

13. Office of the Associate Director for Communication DMB, Division of Public Affairs. October 14, 2016 2016. Seasonal Influenza Vaccine Effectiveness, 2005-2016. Accessed February 22.

14. Castilla J, Godoy P, Domínguez A, Martínez-Baz I, Astray J, Martín V, Delgado-Rodríguez M, Baricot M, Soldevila N, Mayoral JM, Quintana JM, Galán JC, Castro A, González-Candelas F, Garín O, Saez M, Tamames S, Pumarola T, Spain CCaCiIWG. 2013. Influenza vaccine effectiveness in preventing outpatient, inpatient, and severe cases of laboratory-confirmed influenza. Clin Infect Dis 57:167–75.

15. Puig-Barberà J, Arnedo-Pena A, Pardo-Serrano F, Tirado-Balaguer MD, Pérez-Vilar S, Silvestre-Silvestre E, Calvo-Mas C, Safont-Adsuara L, Ruiz-García M, Surveillance and Vaccine Evaluation Group during the autumn 2009 H1N1 pandemic wave in Castellón Sa. 2010. Effectiveness of seasonal 2008-2009, 2009-2010 and pandemic vaccines, to prevent influenza hospitalizations during the autumn 2009 influenza pandemic wave in Castellón, Spain. A test-negative, hospital-based, case-control study. Vaccine 28:7460–7.

16. Szilagyi PG, Fairbrother G, Griffin MR, Hornung RW, Donauer S, Morrow A, Altaye M, Zhu Y, Ambrose S, Edwards KM, Poehling KA, Lofthus G, Holloway M, Finelli L, Iwane M, Staat MA, Network NVS. 2008. Influenza vaccine effectiveness among children 6 to 59 months of age during 2 influenza seasons: a case-cohort study. Arch Pediatr Adolesc Med 162:943–51.

17. Russell KL, Ryan MAK, Hawksworth A, Freed NE, Irvine M, Daum LT. 2005. Effectiveness of the 2003–2004 influenza vaccine among U.S. military basic trainees: a year of suboptimal match between vaccine and circulating strain. 23:1981–1985.

18. Eichelberger MC, Morens DM, Taubenberger JK. 2018. Neuraminidase as an influenza vaccine antigen: a low hanging fruit, ready for picking to improve vaccine effectiveness. 53:38–44.

19. Krammer F, Fouchier RAM, Eichelberger MC, Webby RJ, Shaw-Saliba K, Wan H, Wilson PC, Compans RW, Skountzou I, Monto AS. 2018. NAction! How Can Neuraminidase-Based Immunity Contribute to Better Influenza Virus Vaccines? mBio 9.

20. Krammer F, Li L, Wilson PC. 2019. Emerging from the Shadow of Hemagglutinin: Neuraminidase Is an Important Target for Influenza Vaccination. 26:712–713.

21. Jagadesh A, Salam AAA, Mudgal PP, Arunkumar G. 2016. Influenza virus neuraminidase (NA): a target for antivirals and vaccines. 161:2087–2094.

22. Gilchuk IM, Bangaru S, Gilchuk P, Irving RP, Kose N, Bombardi RG, Thornburg NJ, Creech CB, Edwards KM, Li S, Turner HL, Yu W, Zhu X, Wilson IA, Ward AB, Crowe JE. 2019. Influenza H7N9 Virus Neuraminidase-Specific Human Monoclonal Antibodies Inhibit Viral Egress and Protect from Lethal Influenza Infection in Mice. 26:715-728.e8.

23. Zhu X, Turner HL, Lang S, McBride R, Bangaru S, Gilchuk IM, Yu W, Paulson JC, Crowe JE, Ward AB, Wilson IA. 2019. Structural Basis of Protection against H7N9 Influenza Virus by Human Anti-N9 Neuraminidase Antibodies. 26:729-738.e4.

24. Maier HE, Nachbagauer R, Kuan G, Ng S, Lopez R, Sanchez N, Stadlbauer D, Gresh L, Schiller A, Rajabhathor A, Ojeda S, Guglia AF, Amanat F, Balmaseda A, Krammer F, Gordon A. 2019. Pre-existing anti-neuraminidase antibodies are associated with shortened duration of influenza A (H1N1)pdm virus shedding and illness in naturally infected adults. Clin Infect Dis.

25. Walz L, Kays SK, Zimmer G, von Messling V. 2018. Neuraminidase-Inhibiting Antibody Titers Correlate with Protection from Heterologous Influenza Virus Strains of the Same Neuraminidase Subtype. J Virol 92.

26. Memoli MJ, Shaw PA, Han A, Czajkowski L, Reed S, Athota R, Bristol T, Fargis S, Risos K, Powers JH, Davey RT, Taubenberger JK. 2016. Evaluation of Antihemagglutinin and Antineuraminidase Antibodies as Correlates of Protection in an Influenza A/H1N1 Virus Healthy Human Challenge Model. mBio 7:e00417–16.

27. Wohlbold TJ, Chromikova V, Tan GS, Meade P, Amanat F, Comella P, Hirsh A, Krammer F. 2015. Hemagglutinin Stalk-and Neuraminidase-Specific Monoclonal Antibodies Protect against Lethal H10N8 Influenza Virus Infection in Mice. J Virol 90:851–61.

28. Wan H, Gao J, Xu K, Chen H, Couzens LK, Rivers KH, Easterbrook JD, Yang K, Zhong L, Rajabi M, Ye J, Sultana I, Wan X-F, Liu X, Perez DR, Taubenberger JK, Eichelberger MC. 2013. Molecular Basis for Broad Neuraminidase Immunity: Conserved Epitopes in Seasonal and Pandemic H1N1 as Well as H5N1 Influenza Viruses. Journal of Virology 87:9290.

29. Wan H, Yang H, Shore DA, Garten RJ, Couzens L, Gao J, Jiang L, Carney PJ, Villanueva J, Stevens J, Eichelberger MC. 2015. Structural characterization of a protective epitope spanning A(H1N1)pdm09 influenza virus neuraminidase monomers. Nature communications 6:6114–6114.

30. Jiang L, Fantoni G, Couzens L, Gao J, Plant E, Ye Z, Eichelberger MC, Wan H. 2015. Comparative Efficacy of Monoclonal Antibodies That Bind to Different Epitopes of the 2009 Pandemic H1N1 Influenza Virus Neuraminidase. Journal of virology 90:117–128.

31. Gao J, Couzens L, Burke DF, Wan H, Wilson P, Memoli MJ, Xu X, Harvey R, Wrammert J, Ahmed R, Taubenberger JK, Smith DJ, Fouchier RAM, Eichelberger MC. 2019. Antigenic Drift of the Influenza A(H1N1)pdm09 Virus Neuraminidase Results in Reduced Effectiveness of A/California/7/2009 (H1N1pdm09)-Specific Antibodies. mBio 10.

32. Yasuhara A, Yamayoshi S, Kiso M, Sakai-Tagawa Y, Koga M, Adachi E, Kikuchi T, Wang IH, Yamada S, Kawaoka Y. 2019. Antigenic drift originating from changes to the lateral surface of the neuraminidase head of influenza A virus. 4:1024–1034.

33. Stadlbauer D, Zhu X, McMahon M, Turner JS, Wohlbold TJ, Schmitz AJ, Strohmeier S, Yu W, Nachbagauer R, Mudd PA, Wilson IA, Ellebedy AH, Krammer F. 2019. Broadly protective human antibodies that target the active site of influenza virus neuraminidase. Science 366:499.

34. Duehr J, Wohlbold TJ, Oestereich L, Chromikova V, Amanat F, Rajendran M, Gomez-Medina S, Mena I, tenOever BR, Garcia-Sastre A, Basler CF, Munoz-Fontela C, Krammer F. 2017. Novel Cross-Reactive Monoclonal Antibodies against Ebolavirus Glycoproteins Show Protection in a Murine Challenge Model. J Virol 91.

35. Kirkpatrick E, Carreno JM, McMahon M, Jiang K, van Bakel H, Wilson P, Krammer F. 2021. Mutations in the HA stalk domain do not permit escape from a protective, stalk-based vaccine induced immune response in the mouse model. mBio In press.

36. Tan GS, Krammer F, Eggink D, Kongchanagul A, Moran TM, Palese P. 2012. A Pan-H1 Anti-Hemagglutinin Monoclonal Antibody with Potent Broad-Spectrum Efficacy In Vivo, p 6179-88, JVirol, vol 86, 1752 N St., N.W., Washington, DC.

37. Mena I, Nelson MI, Quezada-Monroy F, Dutta J, Cortes-Fernández R, Lara-Puente JH, Castro-Peralta F, Cunha LF, Trovão NS, Lozano-Dubernard B, Rambaut A, van Bakel H, García-Sastre A. 2016. Origins of the 2009 H1N1 influenza pandemic in swine in Mexico. Elife 5.

38. Li Q, Qi J, Zhang W, Vavricka CJ, Shi Y, Wei J, Feng E, Shen J, Chen J, Liu D, He J, Yan J, Liu H, Jiang H, Teng M, Li X, Gao GF. 2010. The 2009 pandemic H1N1 neuraminidase N1 lacks the 150-cavity in its active site. 17:1266–1268.

